# A curated benchmark of enhancer-gene interactions for evaluating enhancer-target gene prediction methods

**DOI:** 10.1101/745844

**Authors:** Jill E. Moore, Henry Pratt, Michael Purcaro, Zhiping Weng

## Abstract

Many genome-wide collections of candidate cis-regulatory elements (cCREs) have been defined using genomic and epigenomic data, but it remains a major challenge to connect these elements to their target genes. To facilitate the development of computational methods for predicting target genes, we developed a Benchmark of candidate <u>En</u>hancer-<u>G</u>ene Interactions (BENGI) by integrating the Registry of cCREs we developed recently with experimentally-derived genomic interactions. We used BENGI to test several published computational methods for linking enhancers with genes, including signal correlation and the supervised learning methods TargetFinder and PEP. We found that while TargetFinder was the best performing method, it was modestly better than a baseline distance method for most benchmark datasets while trained and tested within the same cell type and that TargetFinder often did not outperform the distance method when applied across cell types. Our results suggest that current computational methods need to be improved and that BENGI presents a useful framework for method development and testing.

## INTRODUCTION

With the rapid increase of genomic and epigenomic data in recent years, our ability to annotate regulatory elements across the human genome and predicting their activities in specific cell and tissue types has substantially improved. Widely used approaches integrate multiple epigenetic signals, such as chromatin accessibility, histone marks, and transcribed RNAs ^1-7^ to define collections of regulatory elements, which can be used to study the regulatory programs in diverse cell types and dissect the genetic variations associated with human diseases^5,8-11^.

To maximize the utility of regulatory elements, one must know which genes they regulate. We recently developed the Registry of candidate cis-Regulatory elements (cCREs), a collection of candidate regulatory genomic regions in human and mouse, by integrating chromatin accessibility (DNase-seq) data and histone mark ChIP-seq data in hundreds of biosamples generated by the ENCODE Consortium (http://encodeproject.org/SCREEN). Over 75% of these cCREs have enhancer-like signatures (high chromatin accessibility as measured by high DNase-seq signal and high level of the enhancer-specific histone mark H3K27ac) and are distal (> 2 kb) from an annotated transcription start site (TSS). For cCREs proximal to a TSS, it may be safe to assume that the TSS corresponds to the target gene, but to annotate the biological function of the TSS-distal cCREs and interpret the genetic variants they harbor, we need to determine the genes they regulate.

Assigning enhancers to target genes on a genome-wide scale remains a difficult task. While one could assign an enhancer to the closest gene using linear distance, there are many examples of enhancers skipping over nearby genes in favor of more distal targets^12^. Experimental assays such as Hi-C and ChIA-PET survey physical interactions between genomic regions^13,14^ and by overlapping the anchors of these interactions with annotated enhancers and promoters, we can infer regulatory connections. Approaches based on quantitative trait loci (QTL) associate genetic variants in intergenic regions with genes by the variation of their expression levels across multiple individuals in a human population^15,16^. Recently a single-cell perturbation approach extended this idea^17^. However, these assays are expensive to perform and have only been conducted in high resolution in a small number of cell types. Therefore, we need to rely on computational methods to broadly predict enhancer-gene interactions.

One popular computational method for identifying enhancer-gene interactions is correlating genomic and epigenomic signals at enhancers and gene promoters across multiple biosamples. This method is based on the assumption that enhancers and genes tend to be active or inactive in the same cell types. The first study to utilize this method linked enhancers with genes by correlating active histone mark signal at enhancers with gene expression across nine cell types^1^. Several groups subsequently used similar approaches to link enhancers and genes by correlating various combinations of DNase, histone mark, transcription factor, and gene expression data^8,18-20^. While these methods successfully identified a subset of biologically relevant interactions, their performance has yet to be systematically evaluated.

Other groups have developed supervised machine-learning methods that train statistical models on sets of known enhancer-gene pairs. Most of these models use epigenomic signals (e.g., histone marks, TFs, DNase) at the enhancers, promoters, or intervening windows as input features^21-24^. PEP-motif, on the other hand, uses sequence-based features^25^. The performance of these methods has also not been systematically evaluated for several reasons. First, different methods use different definitions for enhancers ranging from EP300 peaks^23^ to chromatin segmentations^24^. Second, these methods use different datasets to define their gold standards, such as ChIA-PET interactions^21,23^ or Hi-C loops^23,24^, along with different methods for generating negative pairs. Finally, many of these methods use a traditional randomized cross-validation scheme, which results in severe overfitting of some supervised models due to overlapping features^26,27^

To facilitate the development of target gene-prediction methods, we developed a collection of benchmark datasets by integrating the Registry of cCREs with experimentally-derived genomic interactions. We then tested several published methods for linking enhancers with genes, including signal correlation and the supervised learning methods TargetFinder and PEP^24,25^. Overall, we found that while TargetFinder was the best performing method, it was modestly better than a baseline distance method for most benchmark datasets while trained and tested within the same cell type, and Target Finder often did not outperform the distance method when applied across cell types. Our results suggest that current computational methods need to be improved and that our benchmark presents a useful framework for method development and testing.

## RESULTS

### A <u>B</u>enchmark of candidate <u>En</u>hancer-<u>G</u>ene <u>I</u>nteractions (BENGI)

To effectively evaluate target-gene prediction methods, we curated a <u>B</u>enchmark of candidate <u>En</u>hancer-<u>G</u>ene Interactions (BENGI) by integrating our predicted enhancers, cCREs-ELS, with 3D chromatin interactions, genetic interactions, and CRISPR/dCAS9 perturbations, in total 21 datasets across thirteen biosamples (**Figure 1a, Supplemental Tables 1 and 2a**). For 3D chromatin interactions, which include ChIA-PET, Hi-C, and CHi-C datasets, we selected all links with one anchor overlapping a cCRE-ELS and the other anchor falling within 2 kb of a GENCODE annotated TSS (**Figure 1b**, *see Methods*). For about three-quarters of the interactions in total, the anchor of the 3D chromatin interaction overlaps the proximal region of more than one gene making the assignment of the exact gene target ambiguous. To assess the impact of these potentially ambiguous assignments, we created two versions of each 3D interaction benchmark dataset. In the first, we retained all cCRE-gene links; in the second, we removed links with ends within 2 kb of the TSSs of multiple genes (i.e., ambiguous pairs). For genetic interactions (eQTLs) and CRISPR/dCas9 perturbations (crisprQTLs), we paired a cCRE-ELS with a gene if the cCRE overlapped the reported SNP or targeted region (**Figure 1b**). In total, we curated over 185 thousand unique cCRE-gene pairs across the thirteen biosamples. Because these experimental datasets capture different aspects of enhancer-gene interactions (see statistical analyses in the next section), we keep the cCRE-gene pairs as separate datasets in BENGI.

**Figure 1.**
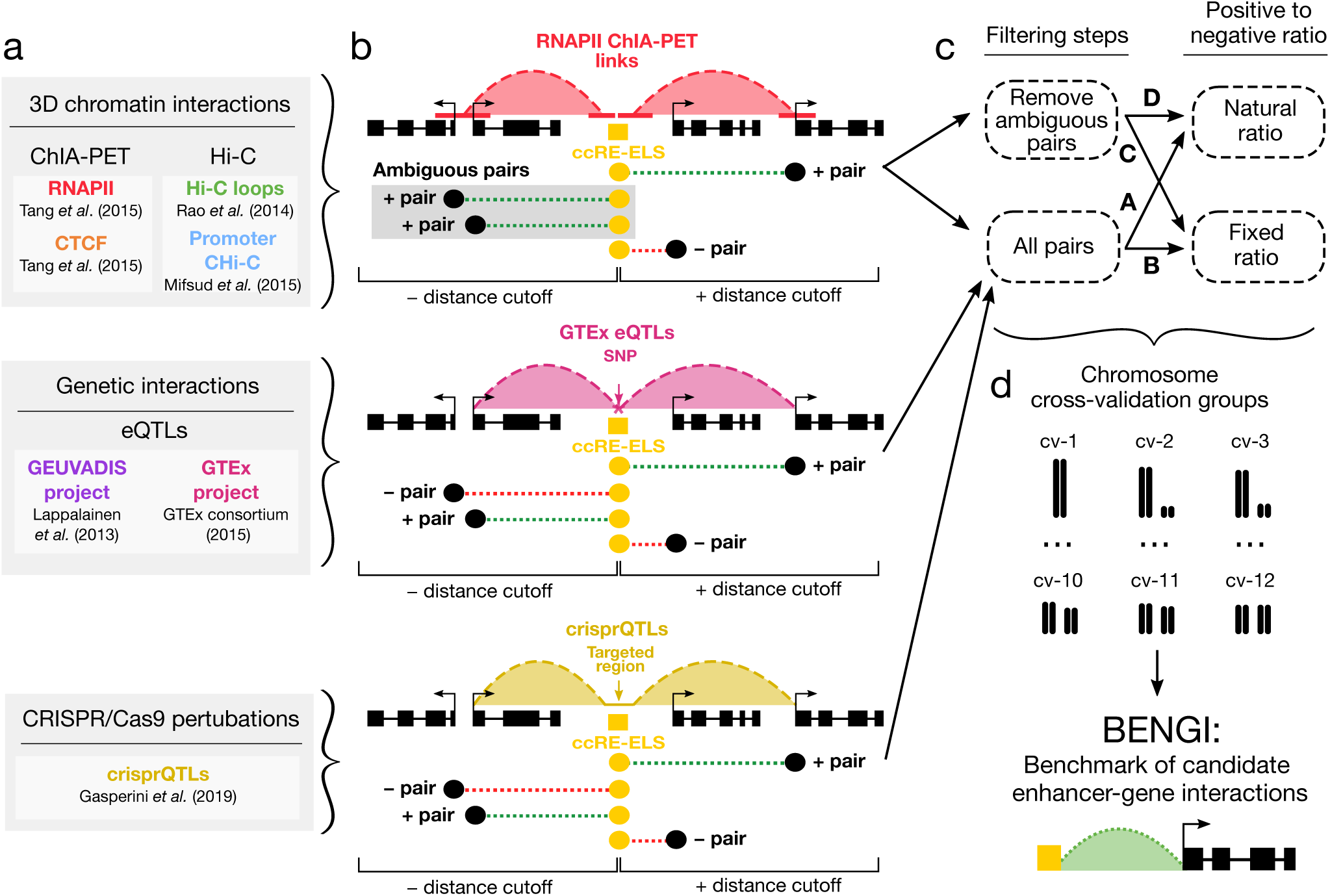
A benchmark of candidate enhancer-gene interactions (BENGI). a, Experimental datasets used to curate BENGI interactions categorized by 3D chromatin interactions, genetic interactions, and CRISPR/Cas9 perturbations. **b**, Methods of generating cCRE-gene pairs (dashed straight lines in green, shaded green, or red) from experimentally determined interactions or perturbation links (dashed, shaded arcs in red, pink, or gold). Each cCRE-gene pair derived from 3D chromatin interactions (top panel) has a cCRE-ELS (yellow box) intersecting one anchor of a link, and the pair is classified depending on the other anchor of the link: for a positive pair (green dashed line), the other anchor overlaps one or more TSSs of just one gene; for an ambiguous pair (dashed line with grey shade), the other anchor overlaps the TSSs of multiple genes; while for a negative pair (red dashed line), the other anchor does not overlap a TSS. Each cCRE-gene pair derived from genetic interactions or perturbation links (middle and bottom panels) has a cCRE-ELS (yellow box) intersecting an eQTL SNP or a CRISPR-targeted region, and the pair is classified as positive (green dashed line) if the gene is an eQTL or crisprQTL gene, while all the pairs that this cCRE forms with non-eQTL genes that have a TSS within the distance cutoff are considered negative pairs (red dashed line). **c**, To reduce potential false positives from 3D interaction data, we implemented a filtering step to remove ambiguous pairs (gray box in **b**) that link cCREs-ELS to more than one gene. This filtering step was not required for assays that explicitly list the linked gene (eQTLs and crisprQTLs). Additionally, to compare between BENGI datasets, we also curated matching sets of interactions with a fixed positive-to-negative ratio. Therefore, in total four BENGI datasets are curated for each 3D chromatin experiment (A, B, C, D) and two are curated for each genetic interaction and CRISPR/Cas-9 perturbation experiment (A, B). **d**, To avoid overfitting of machine-learning algorithms, all cCRE-gene pairs were assigned to cross-validation (CV) groups based on their chromosomal locations. Positive and negative pairs in the same chromosome were assigned to the same CV group, and chromosomes with complementary sizes were assigned to the same CV group so that the groups had approximately the same number of pairs.

To complement the positive cCRE-gene pairs in each BENGI dataset, we generated negative pairs for each cCRE-ELS by selecting all unpaired genes with a TSS within (either upstream or downstream) the 95th percentile distance of all positive cCRE-gene pairs in the dataset (**Supplemental Table 2a**, *see Methods*). These distance cutoffs ranged from 119 kb (RNAPII ChIA-PET in HeLa) to 1.77 Mb (Hi-C in K562). The percentages of positives pairs also varied from 1.9% (Hi-C in K562) to 23.8% (CHi-C in GM12878), and datasets with greater class imbalance (i.e., smaller percentage of positives) are inherently more challenging for a computational algorithm. To enable the comparison of algorithm performance across datasets, we further created datasets with a fixed ratio of one positive to four negatives for each BENGI dataset, by randomly discarding the excess negatives. This consideration, along with the previously mentioned removal of ambiguous pairs for 3D chromatin interactions, results in four BENGI datasets per ChIA-PET, Hi-C, or CHi-C experiment and two BENGI datasets per eQTL or crisprQTL experiment (**Figure 1c, Supplemental Table 2a**). All pairs with the natural positive-negative ratio were used in our analyses unless otherwise noted.

To facilitate the training and testing of supervised machine-learning algorithms, we then assigned both positive and negative pairs to 12 cross-validation (CV) groups by chromosome such that pairs within the same chromosome were always assigned to the same CV group while different CV groups maintained similar sizes by pairing one large chromosome with one small chromosome (chromCV, see *Methods*, **Figure 1d**). Because GM12878 and other lymphoblastoid cell lines (LCLs) had the most BENGI datasets and have been extensively surveyed by the ENCODE and 1000 Genomes Consortia, we will highlight our analyses on BENGI datasets from LCLs.

### Summary statistics of BENGI datasets

We asked whether the various chromatin, genetic, and CRISPR experiments might capture different types of enhancer-gene interactions. To answer this question, we carried out several statistical analyses across the BENGI datasets. First, we performed hierarchical clustering on the six BENGI datasets in GM12878/LCLs by overlap coefficient—the number of positive cCRE-gene pairs shared between two datasets divided by the number of positives in the smaller dataset. Two clusters resulted; one comprised the two eQTL datasets and the other the four chromatin interaction datasets (**Figure 2b**). This overall grouping of the datasets is consistent with the characteristics of the experimental techniques (**Table 1**). Beyond the overall grouping, the two eQTL datasets had higher overlap coefficients with the RNAPII ChIA-PET and CHi-C datasets (0.20–0.36) than with the Hi-C and CTCF ChIA-PET datasets (0.01–0.05). This reflects the promoter emphasis of the first four techniques, hence enriching for promoter-proximal interactions. In contrast, Hi-C identifies significantly more distant interactions than the other techniques (**Figure 2a**, Wilcoxon rank-sum test *p*-value = 1.1E-223).

**Table 1.**
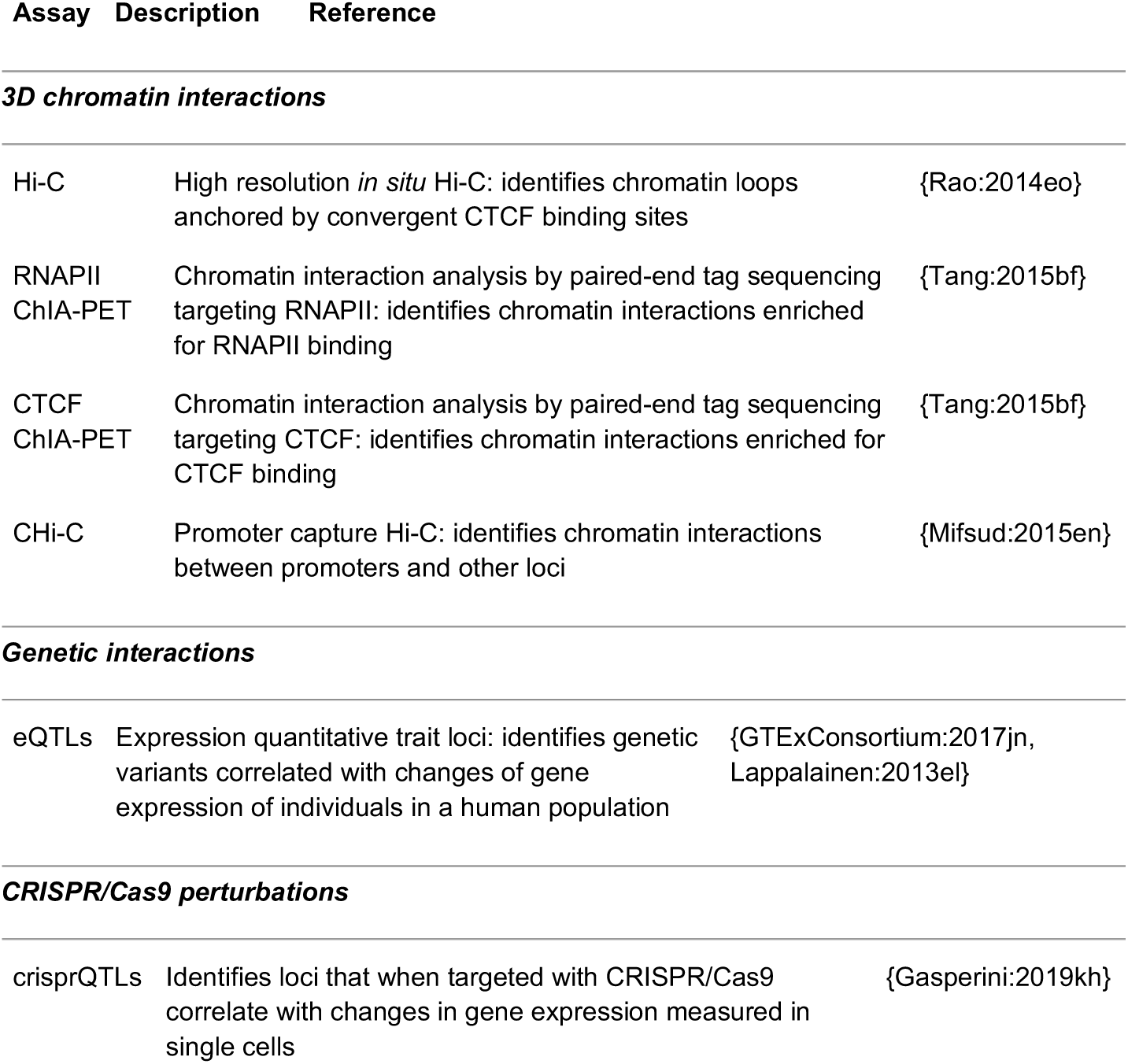
Genomic interaction datasets.

| **Assay** | **Description** | **Reference** |
| --- | --- | --- |
| ***3D chromatin interactions*** |  |  |
| Hi-C | High resolution *in situ* Hi-C: identifies chromatin loops anchored by convergent CTCF binding sites | {Rao:2014eo} |
| RNAPII ChIA-PET | Chromatin interaction analysis by paired-end tag sequencing targeting RNAPII: identifies chromatin interactions enriched for RNAPII binding | {Tang:2015bf} |
| CTCF ChIA-PET | Chromatin interaction analysis by paired-end tag sequencing targeting CTCF: identifies chromatin interactions enriched for CTCF binding | {Tang:2015bf} |
| CHi-C | Promoter capture Hi-C: identifies chromatin interactions between promoters and other loci | {Mifsud:2015en} |
| ***Genetic interactions*** |  |  |
| eQTLs | Expression quantitative trait loci: identifies genetic variants correlated with changes of gene expression of individuals in a human population | {GTExConsortium:2017jn, Lappalainen:2013el} |
| ***CRISPR/Cas9 perturbations*** |  |  |
| crisprQTLs | Identifies loci that when targeted with CRISPR/Cas9 correlate with changes in gene expression measured in single cells | {Gasperini:2019kh} |

**Figure 2.**
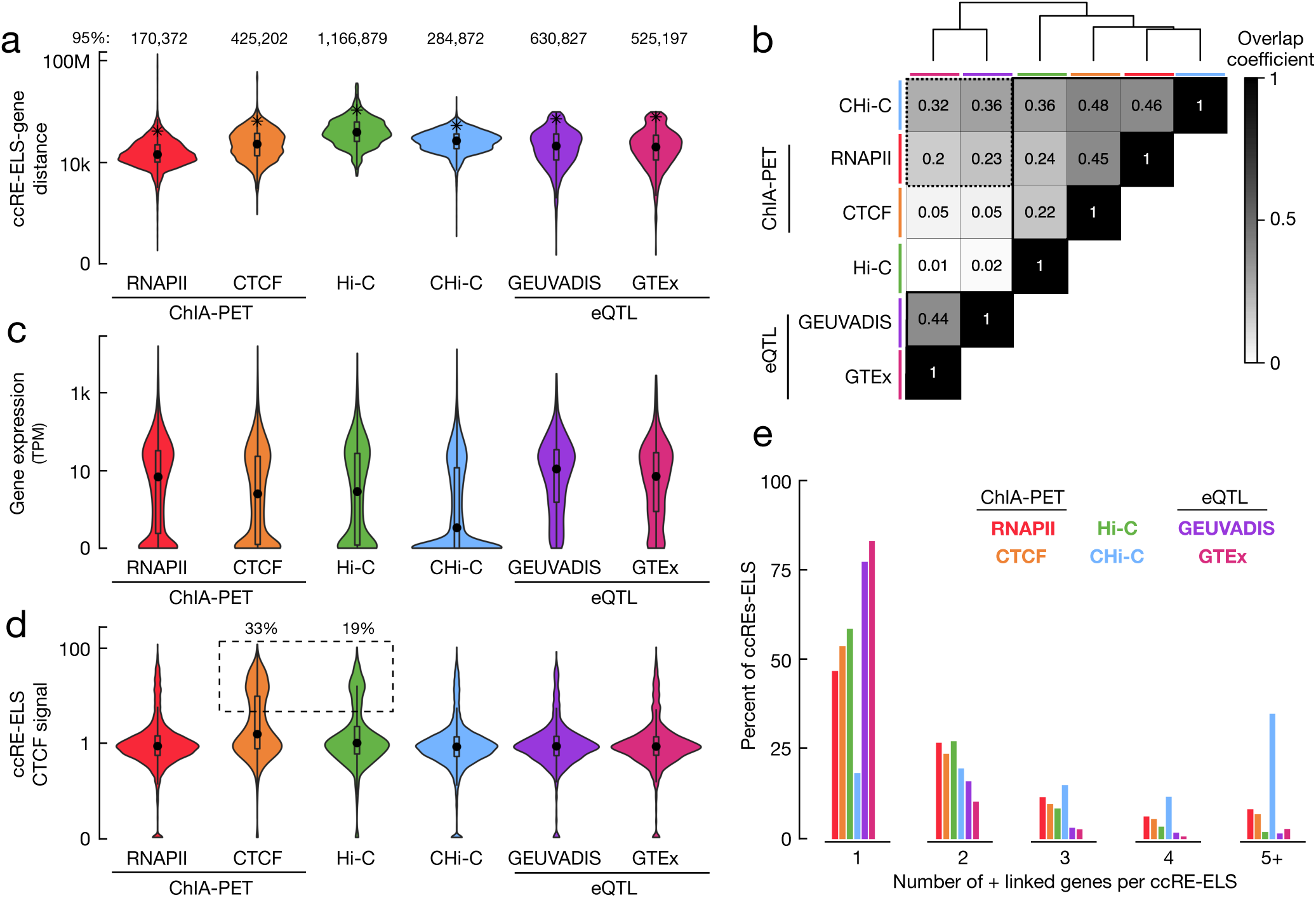
Characteristics of BENGI datasets. Six datasets in GM12878 or other LCLs were evaluated: RNAPII ChIA-PET (red), CTCF ChIA-PET (orange), Hi-C (green), CHi-C (blue), GEUVADIS eQTLs (purple), and GTEx eQTLs (pink), and the same coloring scheme is used for all panels. **a**, Violin plots depicting the distance distributions of positive cCRE-gene pairs for each BENGI dataset. The 95th percentile of each distribution is indicated by a star and stated above each plot. **b**, Heatmap depicting the overlap coefficients between positive cCRE-gene pairs in each BENGI dataset. The datasets were clustered using the hclust algorithm and clustered datasets are outlined in black. **c**, Violin plots depicting the expression levels of genes in positive cCRE-gene pairs (in transcripts per million, TPM). **d**, Violin plots depicting CTCF signal levels at cCREs-ELSs in positive cCRE-gene pairs. A dashed box points out cCREs-ELS with signal > 5. **e**, Distributions of the number of genes positively linked with a cCRE-ELS across datasets.

We then compared gene expression of the positive pairs among the six GM12878/LCL datasets (**Figure 2c**). Overall, genes in the GEUVADIS eQTLs pairs had the highest median expression (median = 10.9 transcripts per million sequenced reads or TPM, Wilcoxon rank-sum test *p* = 1E-3), while genes in the CHi-C pairs had the lowest median expression levels (median = 0.24 TPM, *p* = 7E-39). When we removed ambiguous pairs, gene expression increased significantly for all four chromatin interaction datasets (**Supplemental Figure 1a**), suggesting that some of the ambiguous pairs are false positives. We observed similar increases in gene expression upon removal of ambiguous pairs in other cell types for which we had RNA-seq data (**Supplemental Figure 1b-d**). Without ambiguous pairs, RNAPII ChIA-PET pairs have comparable expression to GEUVADIS eQTL pairs. The enrichment for RNAPII in the ChIA-PET protocol may preferentially identify interactions that involve higher RNAPII activity and higher gene expression. The K562 crisprQTL pairs had the highest overall median expression of 26.4 TPM. We expected to see high expression for the eQTL and crisprQTL datasets because these interactions can only be detected for genes that are expressed in their respective biosamples.

We also observed significant differences in CTCF ChIP-seq signals at cCREs-ELS between BENGI datasets—cCREs-ELS in CTCF ChIA-PET pairs and Hi-C pairs showed significantly higher CTCF signals than cCREs-ELS in other datasets (Wilcoxon rank-sum test *p* < 3.7E-9, **Figure 2d, Supplemental Table 2b)**. Similarly, these pairs were enriched for components of the cohesin complex such as RAD21 and SMC3 (**Supplemental Table 2b**). This enrichment for CTCF is biologically consistent as CTCF was the target for the ChIA-PET experiment, and Hi-C loops are enriched for convergent CTCF binding sites^28^.

Finally, we tallied the number of linked genes for each cCRE-ELS. Across all BENGI datasets, the majority of cCREs-ELS were linked to just one target gene (**Figure 2e, Supplemental Table 2c**). As expected, this trend was more pronounced for 3D chromatin datasets without ambiguous pairs (on average, 84% of cCREs-ELS were paired with only one gene, *p* < 3.3E-5). With or without ambiguous pairs, a lower percentage of cCREs-ELS in CHi-C pairs was paired with just one gene (18% of all pairs and 55% of unambiguous pairs) than in other BENGI datasets (*p* < 3.1E-75). This observation, along with the lower average expression of the linked genes (**Figure 2c**), suggests that either some of the CHi-C pairs were false positives or they captured interactions between cCREs-ELS and genes yet to be expressed.

These analyses suggest that the various experimental techniques, the results of which formed the basis of the BENGI datasets, capture different classes of genomic interactions. Because we do not have a complete understanding of which experimental techniques was best able to capture bona fide enhancer-gene interactions, we propose that computational methods (**Table 2**) should be evaluated on the entire collection of these BENGI datasets to provide a comprehensive understanding of their performance.

**Table 2.**
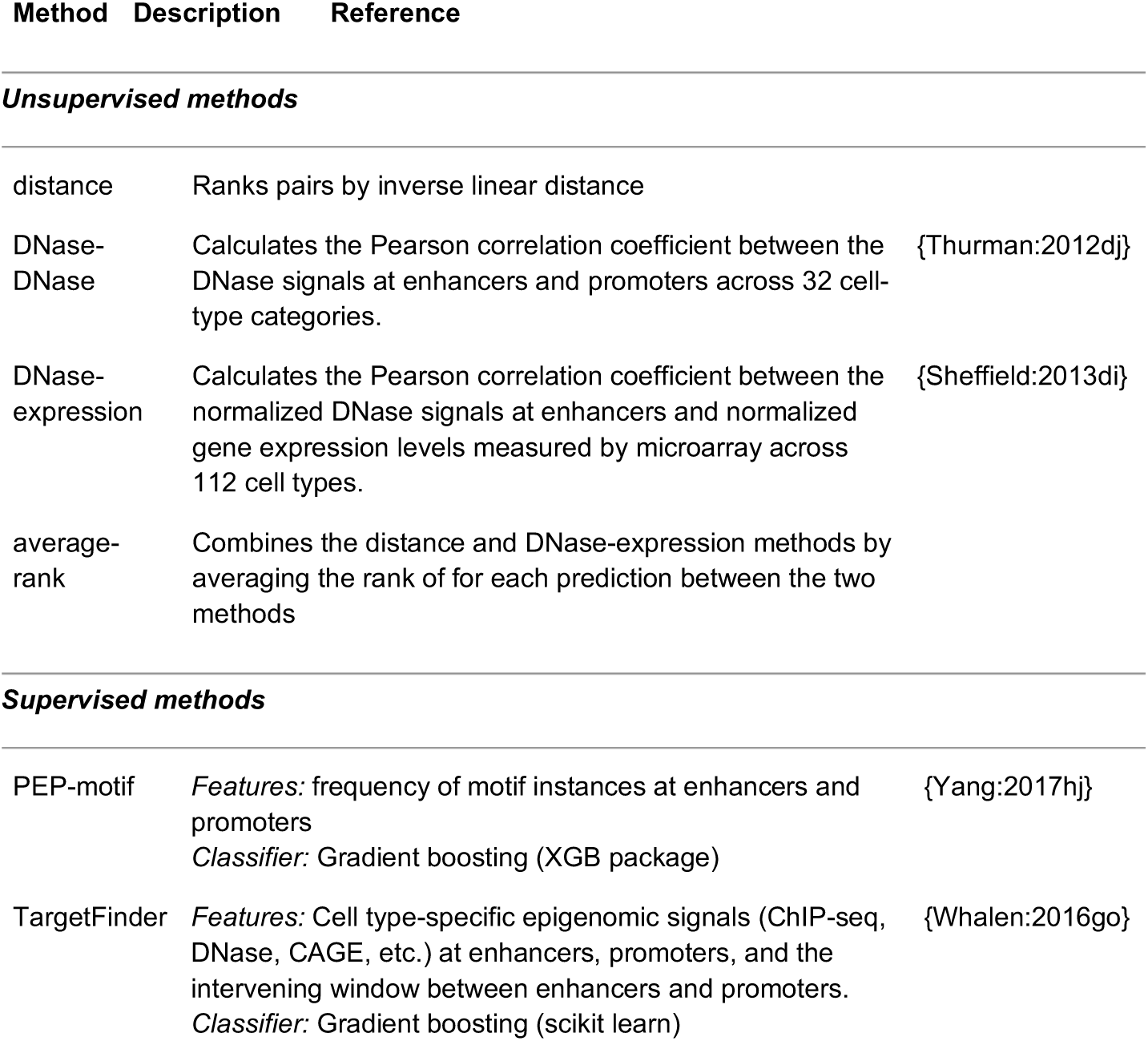
Computational methods for target gene prediction.

### A baseline method of target-gene prediction using genomic distance

Using the BENGI datasets, we evaluated a simple *closest-gene* method for target-gene prediction: assigning a cCRE-ELS to its closest gene in linear distance, computed by subtracting genomic coordinates of the cCRE and the nearest TSS. We tested this method using two gene sets—all genes or all protein-coding genes annotated by GENCODE V19—by evaluating precision and recall on each BENGI dataset. Using protein-coding genes performed invariably better than using all genes (on average 50% better over all 21 datasets across cell types; **Supplemental Table 2d);** thus, we used protein-coding genes for all subsequent analyses with this method.

The *closest-gene* method worked best for crisprQTL pairs (precision = 0.68 and recall = 0.62) followed by ChIA-PET RNAPII pairs (precision = 0.60 and recall = 0.33 averaged across cell lines). The method performed the worst for Hi-C pairs with an average precision of 0.18 and an average recall of 0.11. These results are consistent with our statistical analyses described above, which revealed that crisprQTL and RNAPII ChIA-PET pairs were enriched in gene-proximal interactions while Hi-C pairs tended to identify more distal interactions.

To compare with other enhancer-gene prediction methods, we adapted the *closest-gene* method to a quantitative ranking scheme where we ordered cCRE-gene pairs by the distance between the cCRE-ELS and the gene’s closest TSS. For each BENGI dataset, we evaluated the overall performance of the resulting *distance* method by calculating the area under the precision-recall curve (AUPR). Accordingly, the *distance* method had the highest AUPR (0.47) for crisprQTL pairs and the lowest AUPR (0.06) for Hi-C pairs (**Figure 3a,b, Supplemental Figure 2b, Supplemental Table 3**). Since the distance method is cell-type independent and does not require any experimental data, we considered it the baseline method for comparing all enhancer-gene prediction methods.

**Figure 3.**
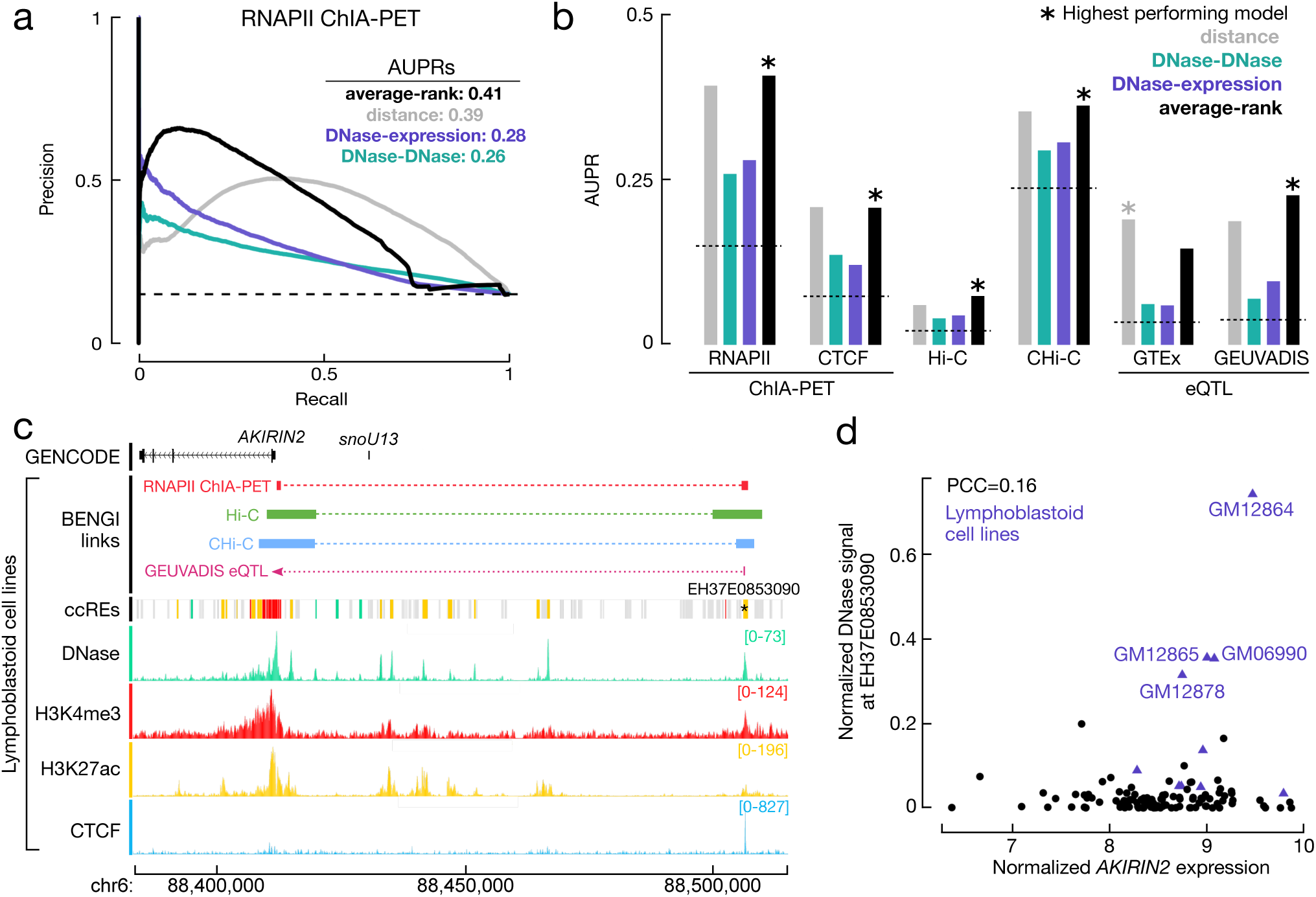
Evaluating unsupervised methods for predicting cCRE-gene pairs. **a**, Precision-recall (PR) curves for four unsupervised methods evaluated on RNAPII ChIA-PET pairs in GM12878: the distance between cCREs-ELS and genes (gray), DNase-DNase correlation by Thurman *et al.* (green), DNase-expression correlation by Sheffield *et al.* (purple), and the average rank of distance and the DNase-expression method (black). Areas under the PR curve (AUPRs) for the four methods are listed in the legend. The AUPR for a random method is indicated with the dashed line at 0.14. **b**, AUPRs for the four unsupervised methods in **a** computed on each of the six benchmark datasets in LCLs. **c**, Genome browser view (chr6:88,382,922-88,515,031) of epigenomic signals and positive BENGI links (RNAPII ChIA-PET in red, Hi-C in green, CHi-C in blue, and GEUVADIS eQTL in pink) connecting the cCRE EH37E0853090 (star) to the gene *AKIRIN2*. **d**, Scatter plot of normalized *AKIRIN2* expression vs. normalized DNase signal at EH37E0853090 as calculated by Sheffield et al. (Pearson correlation coefficient = 0.16). Even though *AKIRIN2* is highly expressed across many tissues, EH37E0853090 has high DNase signals primarily in lymphoblastoid cell lines (purple triangles), resulting in a low correlation.

### Correlation-based approaches performed worse than the distance method

We next evaluated the performance of two correlation-based methods on the BENGI datasets: a method based on correlating the DNase signals at predicted enhancers with the DNase signals at TSSs across a panel of biosamples^19^ and a method based on correlating DNase signal with gene expression^20^. Both DNase-DNase and DNase-expression methods outperformed random predictions for all 21 BENGI datasets, with the average AUPR of 0.10 and 0.12 vs. 0.07, respectively, but the differences were modest (**Supplemental Figure 1**; **Supplemental Table 3**). As previously demonstrated^19^, positive pairs had significantly higher correlations for both methods than negative pairs in all datasets (**Supplemental Figure 2**); however, the relative rankings of these correlations were mixed and did not segregate positive from negative pairs completely. The DNase-expression method significantly outperformed the DNase-DNase method in all but two BENGI datasets (Wilcoxon signed-rank test p = 1.0E-4), with an average AUPR increase of 25% (**Supplemental Table 2**).

However, the distance method substantially outperformed these two correlation-based methods: distance was better than DNase-DNase for all 21 datasets (average AUPR increase of 128%; *p* = 9.5E-7; **Supplemental Table 2**) and better than DNase-expression for 17 datasets (average AUPR increase of 82%; *p* = 1.6E-4). The PR curves of the distance and two correlation-based methods on RNAPII ChIA-PET pairs are shown in **Figure 3a**. For the first 25 k predictions, the distance method had a similar precision to the DNase-DNase method and lower than the DNase-expression method, but with more predictions being made, the distance method substantially outperformed both correlation-based methods and achieved a much higher AUPR (0.39 vs. 0.26 and 0.28). We observed this cross-over of PR curves in other non-QTL datasets as well (**Supplemental Figure 1**); thus, we integrated the distance and DNase-expression methods by averaging their ranks for the same prediction. Notably, this average-rank method showed high precisions for its top-ranked predictions (**Figure 3a**) and achieved higher AUPRs than the other methods for all 13 datasets except GTEx eQTL and crisprQTL, with an average AUPR increase of 17% over the distance method for these datasets (**Figure 3b, Supplemental Table 2**). For the eight GTEx eQTL and crisprQTL datasets, the distance method remained the best, showing on average 23% higher AUPR than the second-best method, average-rank (**Supplemental Table 2**).

We asked why correlation-based methods performed poorly for predicting enhancer-gene pairs. One particular example is highlighted in **Figure 3c-d**. The cCRE-ELS EH37E0853090 is paired with the gene *AKIRIN2* by RNAPII ChIA-PET, Hi-C, CHi-C, and a GEUVADIS eQTL (**Figure 3c**). However, this pair is poorly ranked by both correlation-based methods (correlation coefficient *r* = 0.03 and 0.16 for DNase-DNase and DNase-expression, respectively). *AKIRIN2* was highly expressed in most surveyed cell types (median normalized expression 8.5 vs. background of 4.7 RPKM, **Supplemental Figure 4a)** and its promoter had high DNase signal (signal ≥ 50) for each of the DNase-seq groups (**Supplemental Figure 4b**). EH37E0853090, however, only had high DNase signals in four cell types, which were all lymphoblastoid cell lines, suggesting that this enhancer was primarily active in the B cell lineage. The ubiquitous expression of *AKIRIN2* and the cell-type-specific activity of EH37E0853091 resulted in a low correlation (**Figure 3d, Supplemental Figure 4b**). In general, TSS-overlapping cCREs (cCREs-TSS) are active in many more biosamples than distal cCREs-ELS (median of 92 vs. 46 biosamples, *p* = 3.6E-264, **Supplemental Figure 4c-d**). In summary, because the epigenomic signals at cCREs-ELS are far more cell type specific than the epigenomic signals at TSSs and gene expression profiles, correlation across biosamples is a poor method for detecting enhancer-gene pairs.

### Supervised methods upon cross-validation outperformed baseline methods

We tested two supervised machine-learning methods that were reported to perform well in their original publications: TargetFinder, which uses as input features epigenomic signals such as histone mark ChIP-seq, TF ChIP-seq, and DNase-seq in the corresponding cell type, and PEP-motif, which uses the occurrence of TF sequence motifs as features. Xi *et al*. subsequently revealed that the original implementations of cross-validation (CV) by TargetFinder and PEP-motif allowed assignment of enhancer-gene pairs from the same genomic loci to different CV groups, which led to sharing of training and testing data, overfitting of their models, and inflated performance^26^. Thus, we implemented the chromCV method to ensure that pairs from the same chromosome were always assigned to the same CV group (**Figure 1e**; *Methods*).

We first tested these two supervised methods on the six BENGI datasets in GM12878 because this cell type had a large number of epigenomic datasets that could be used as features to train the methods. Although PEP-motif performed better than random, it under-performed the distance method for all but the Hi-C dataset and was far worse than the average-rank method for all six datasets (**Figure 4a-b**; **Supplemental Table 2b**). In contrast, TargetFinder outperformed the average-rank method for all six datasets, with an average AUPR improvement of 61% (**Figure 4a-b**; **Supplemental Table 2**), but the AUPRs were still low, especially for the Hi-C (0.13) and eQTL datasets (0.19 and 0.25).

**Figure 4.**
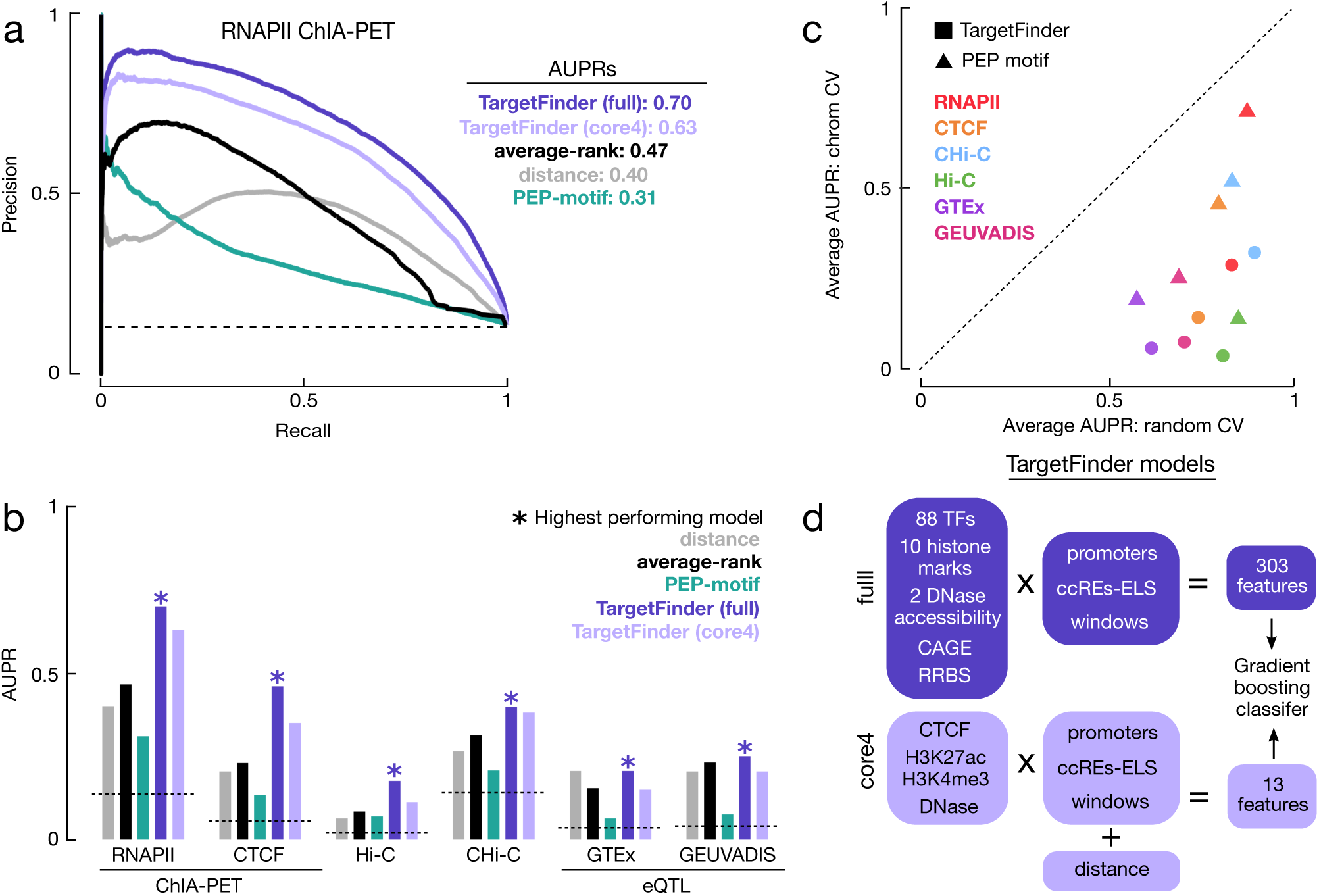
Evaluating supervised learning methods for predicting cCRE-gene pairs. **a**, PR curves for three supervised methods evaluated on RNAPII ChIA-PET pairs in GM12878: PEP-motif (green) and two versions of TargetFinder (full model in darker blue and core model in lighter blue). For comparison, two unsupervised methods from Figure 3, the distance (gray) and average-rank (black) methods, are also shown along with the AUPR for a random method (dashed line at 0.14). AUPRs for the methods are listed in the legend. **b**, AUPR for the three supervised methods, two unsupervised methods, and random, colored as in **a**, for each of the six BENGI datasets in LCLs. **c**, Scatter plot of AUPRs for TargetFinder (squares) and PEP-motif (triangles) across BENGI datasets evaluated using 12-fold random CV (X-axis) vs. chromosome-based CV (Y-axis). The diagonal dashed line indicates X = Y. **d**, Schematic for full and core4 TargetFinder models.

Because the results of TargetFinder and PEP-motif upon our chromCV implementation were worse than their original published results, we also implemented a randomized 12-fold CV method as described in the original publications to test whether we could reproduce their results. Indeed, we observed large performance decreases with the chromCV method with respect to the original CV method (**Figure 4c**), suggesting that overfitting was a source of their inflated performance. PEP-motif had a more substantial drop in performance (average AUPR decrease of 81%) than TargetFinder (average AUPR decrease of 52%), likely because PEP-motif added 4-kb padding on both sides of each enhancer, increasing the chance of overlapping training and testing data. Although PEP-motif and TargetFinder used Hi-C loops as the gold standard in their original analyses, both methods showed the largest performance drops for the BENGI GM12878 Hi-C pairs (AUPR decrease of 96% for PEP-motif and 84% for TargetFinder). This analysis further highlights the utility of a carefully designed benchmark to prevent overfitting of supervised models.

Our implementation of TargetFinder in GM12878 cells used 101 epigenomic datasets, including ChIP-seq of 88 TFs, resulting in a total of 303 input features (**Figure 4d**). However, other biosamples did not have such extensive TF ChIP-seq data; thus, we also trained TargetFinder models using only distance and four epigenomic features—DNase, H3K4me3, H3K27ac, and CTCF data—which we call the core4 TargetFinder models. While the core4 models had an average AUPR reduction of 16% compared with the respective full models across the 13 BENGI datasets (**Figure 4a-b**; **Supplemental Table 3**), they still outperformed the distance and the average-rank methods for all datasets. Of note were IMR-90 Hi-C pairs, which had the greatest drop in performance between the full and core4 TargetFinder models with a reduction of AUPR of 0.28 (80%). We observed similar large decreases in performance across all four variations of IMR-90 Hi-C pairs. Additionally, for GM12878 eQTL and CHiC pairs, the core4 models performed slightly better than the full model, suggesting that the full TargetFinder model may be overfitting these datasets. However, for other variations of these pairs, (fixed ratio and ambiguous pairs removed) the full TargetFinder models had the best performances. We also trained core3 models for the biosamples without CTCF data, and they showed an average AUPR reduction of 28% compared with the respective full models across the 13 BENGI datasets. For the seven GTEx eQTL datasets in tissues, these core3 models did not outperform the distance or average-rank models.

### TargetFinder has a moderate performance cross cell types

The most desirable application of a supervised method is to train the model in a biosample with 3D chromatin or genetic interaction data and then use the model to make predictions in another biosample without such data. Thus, we tested the TargetFinder core4 and core3 models for such an application on the ChIA-PET, Hi-C, CHi-C, and GTEx eQTL datasets readjusting our chromCV to prevent overfitting{Schreiber:ta} (see *Methods*).

As expected, the cross-cell type models performed worse than same-cell type models, but their performance varied when compared with the unsupervised distance and average-rank methods. For CHi-C and RNAPII ChIA-PET datasets, the cross-cell type TargetFinder models outperformed the distance and average-rank methods in both tested cell types (GM12878 vs. HeLa and GM12878 vs. CD34+) with an average AUPR increase of 37% and 12%, respectively (**Figure 5 a,b, Supplemental Table 4**). For CTCF ChIA-PET, the model trained in HeLa did not outperform the unsupervised methods for predicting GM12878 pairs (AUPR = 0.17 vs 0.21 and 0.21) but the model trained in GM12878 did for predicting HeLa pairs (AUPR = 0.26 vs 0.16 and 0.19; **Figure 5c, Supplemental Table 4**). Results for the Hi-C datasets were mixed. Of the 60 cross-cell type models tested, 13 outperformed the distance and average-rank methods. Specifically, the model trained in GM12878 only outperformed distance and average-rank methods for predicting HeLa or NHEK pairs (**Figure 5d, Supplemental Table 4**), with an average 31% increase in performance. The model trained in IMR-90 never outperformed the distance and average-rank methods, and for predicting HMEC, IMR-90, and K562 pairs, none of the cross-cell type models outperformed the distance or average-rank methods (**Supplemental Table 4**). These results were consistent across the fixed-ratio pairs as well. Finally, none of the cross-cell type models outperformed the distance method for GTEx datasets; the distance method was the highest performing model for all GTEx datasets (**Supplemental Table 4**).

**Figure 5.**
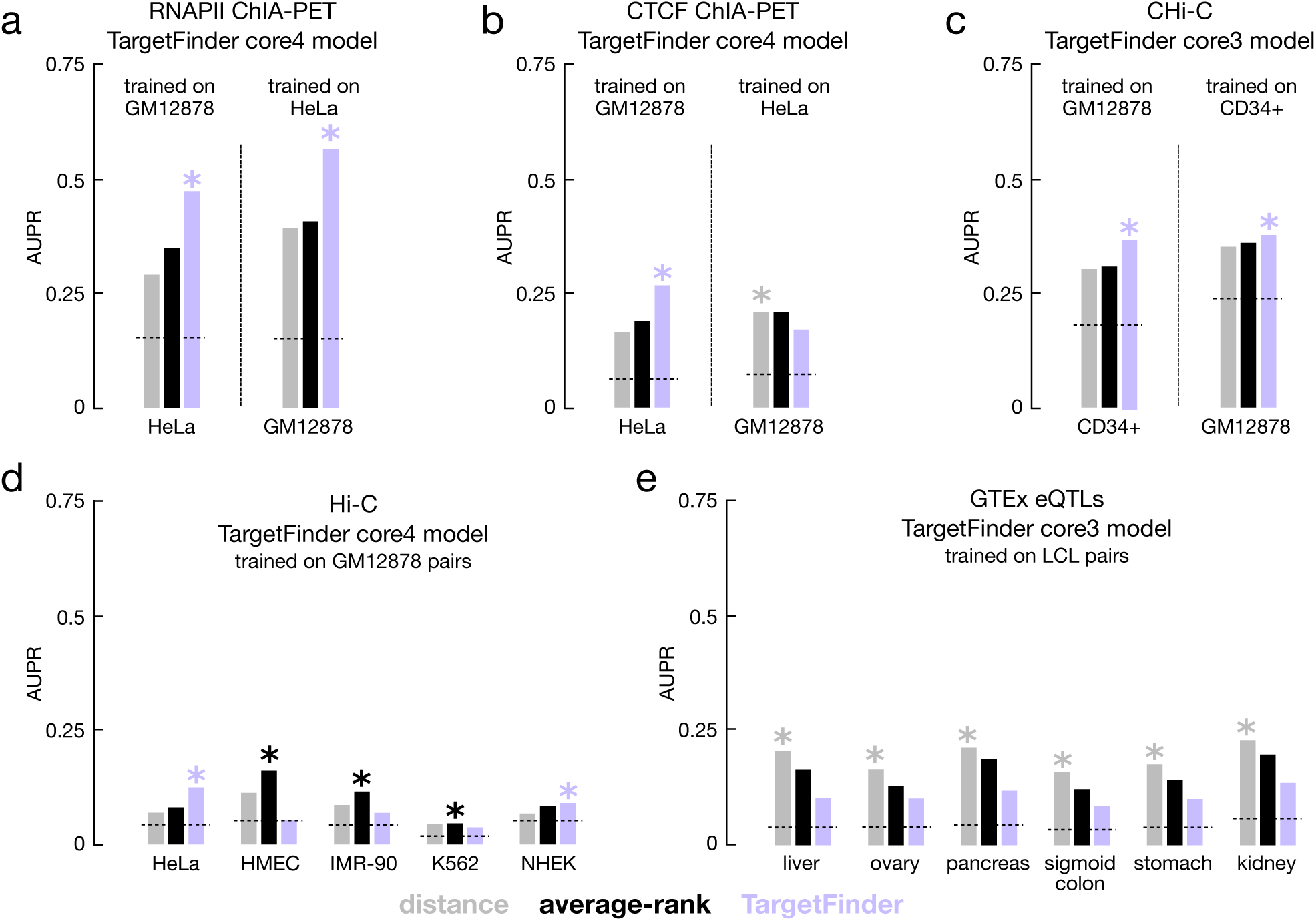
Evaluating supervised learning methods trained in one cell type and tested in another cell type. AUPRs for distance (gray), average-rank (black) and TargetFinder core4 (purple) methods across **a**, RNAPII ChIA-PET **b**, CTCF ChIA-PET **c**, CHi-C **d**, Hi-C, and **e**, GTEx eQTLs pairs. Cell type for training is indicated in the panel title and cell type for testing is indication on the x axis. The best performing method for each dataset is indicated by a star and random performance is indicated with a dashed line.

## DISCUSSION

Here we presented BENGI, a benchmark of cCRE-ELS–gene pairs, curated through the integration of the Registry of cCREs and genomic interaction datasets. We used BENGI to evaluate four published computational methods for target gene prediction that represent most of the widely used approaches in the field while surveying orthogonal dimensions—correlation methods survey across the biosample dimension while supervised machine-learning methods like TargetFinder survey across the assay dimension. We found that the two correlation-based, unsupervised methods significantly underperformed the baseline distance method while one of the two supervised methods examined, TargetFinder, significantly outperformed the distance method when trained and tested within the same cell type by cross-validation. Although TargetFinder outperformed the distance method for all BENGI datasets, the AUPRs of the TargetFinder models were generally still low (0.07-0.71). In particular, TargetFinder performed best on ChIA-PET RNAPII pairs, suggesting that these interactions may be most highly correlated with epigenomic features. The other supervised method, PEP-motif, significantly underperformed the distance method, suggesting that the frequencies of TF motifs at enhancers and promoters are not sufficiently predictive of genomic interactions. When trained and tested in different cell types, TargetFinder beat the distance method for some BENGI datasets, albeit by a much smaller amount. Overall, there is much room for improvement for all of these methods, indicating that target-gene prediction remains a challenging problem. BENGI datasets can be used by the community to tackle this problem while avoiding overfitting issues such as those identified for TargetFinder and PEP post-publication^26,27^.

Our analyses highlight the differences between the genomic interactions as identified by the various experimental techniques (**Table 1**). In the same biosample (e.g., LCLs), the BENGI datasets by the same technique share ∼40% of their pairs (e.g., between RNAPII and CTCF ChIA-PET and between GEUVADIS and GTEx eQTLs), but the overlaps between datasets by different techniques are typically lower than 25% and can be as low as 1% (e.g., between eQTL and Hi-C). The BENGI datasets also differ significantly in enhancer-gene distance and enrichment of epigenomic signals at the enhancers and TSSs. Thus, we still do not have a comprehensive understanding of the factors that regulate enhancer-gene interactions, and the different experimental techniques may be capturing different subsets of interactions.

Overall, all computational methods evaluated had difficulty predicting Hi-C pairs; even in the fixed ratio datasets, the Hi-C pairs consistently had the lowest overall performance. This could be due to the technical challenges of calling Hi-C loops or the biological roles of these loops. For example, it was noted that the detection of Hi-C loops requires care, and different loop calling methods can produce markedly different results^29^. Additionally, recent results from the Aiden lab demonstrated that gene expression did not change upon loop disruption via knocking out the key protein CTCF using a degron system^30^. This may suggest that these CTCF Hi-C loops may serve specific biological roles and may only represent a small subset of enhancer-gene interactions that have different properties compared to the other interactions.

Although the correlation-based methods did not outperform the distance method, the DNase-expression method did augment the distance method when combined with it. Furthermore, because correlation-based methods and supervised machine-learning methods survey orthogonal dimensions (biosample vs. assay), one promising future direction is to combine these two types of approaches. For such future directions to be fruitful, it would be beneficial to understand the different performance between the two correlation-based methods as the DNase-expression correlation method consistently outperformed the DNase-DNase correlation method. Several factors could be contributing to this increased performance. First, gene expression may be a better readout for enhancer-gene interactions than a promoter’s chromatin accessibility, although these two features are correlated (average Pearson correlation *r* = 0.68). Second, for the DNase-expression method, Sheffield *et al.* generated normalized, batch-corrected matrices for the DNase-seq and gene expression data while the DNase-DNase method used read depth normalized signal without any additional processing. To avoid imprecision of reimplementation, we downloaded these exact input datasets from the original publications (i.e., the exact normalized matrices for the DNase-expression method and the ENCODE2-processed DNase-seq bigWigs for the DNase-DNase method). The Sheffield *et al*. normalization technique may correct for outliers and batch effects, which otherwise would lead to spurious correlations impacting performance. Third, the DNase-DNase method merges 79 cell types into 32 groups based on cell type similarity. While this grouping may correct an uneven survey of the biosample space, it may lead to lower overall correlations for cell type-specific interactions. We highlighted one such case with the LCL-specific EH37E0853090-AKIRIN2 interaction where the DNase-DNase method reported a correlation of 0.03, and the DNase-expression method reported a correlation of 0.12. The low correlation by the DNase-DNase method was due to the combination of the four LCLs into one group, reducing statistical power (**Supplemental Figure 4b**). These possible explanations should be carefully considered when designing future correlation-based and combined methods. Additionally, although these correlation-based methods did not perform well on BENGI datasets, they may have better predictive power when used on curated sets of biosamples such as those across embryonic development or cell differentiation. As we expand the number of cell types and tissues covered by BENGI, we hope to test these methods to evaluate their performance systematically.

Finally, we developed BENGI using an enhancer centric model as we were motivated by the Registry of cCREs. We hope to expand upon this to include a gene centric model (i.e., given a gene, determine its interacting enhancers) for future developments. We also plan on expanding BENGI to include more functionally tested datasets such as the crisprQTLs as these results are published. Developing precise and accurate enhancer-gene prediction models will improve our understanding of how regulatory elements control gene expression and ultimately their role in human diseases.

## SUPPLEMENTAL FIGURE CAPTIONS

**Supplemental Figure 1.**
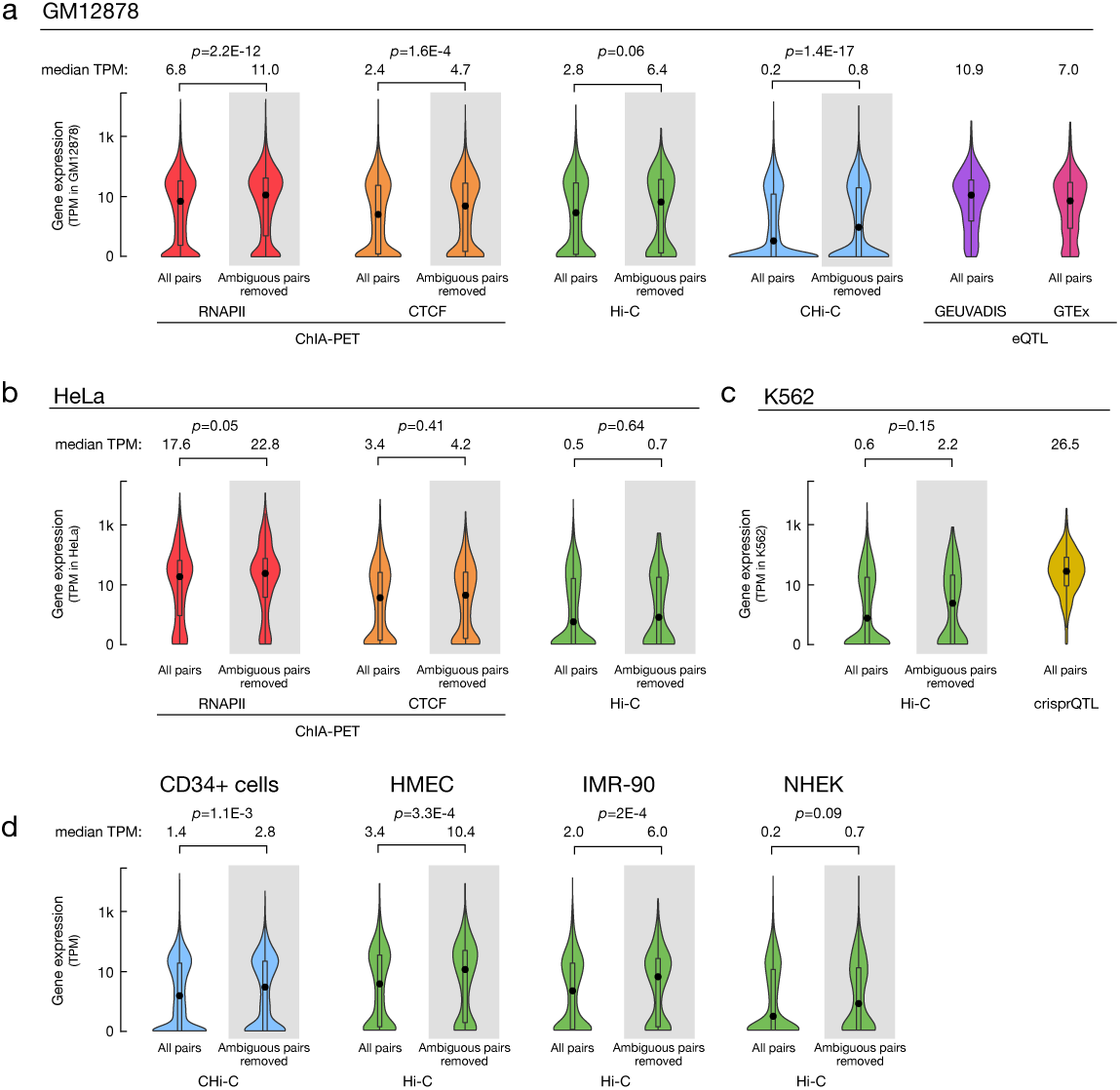
Expression levels of genes in BENGI pairs. Violin plots displaying the distributions of gene expression in positive pairs for each BENGI dataset in **a**, GM12878/LCLs **b**, HeLa **c**, K562, **d**, CD34+ cells, HMEC, IMR-90, and NHEK. The median expression level (in TPM) is displayed above each violin plot. For 3D chromain datasets (ChIA-PET, Hi-C and CHi-C), genes in all positive pairs and with ambiguous pairs removed were compared and Wilcoxon rank-sum test *p*-values are indicated.

**Supplemental Figure 2.**
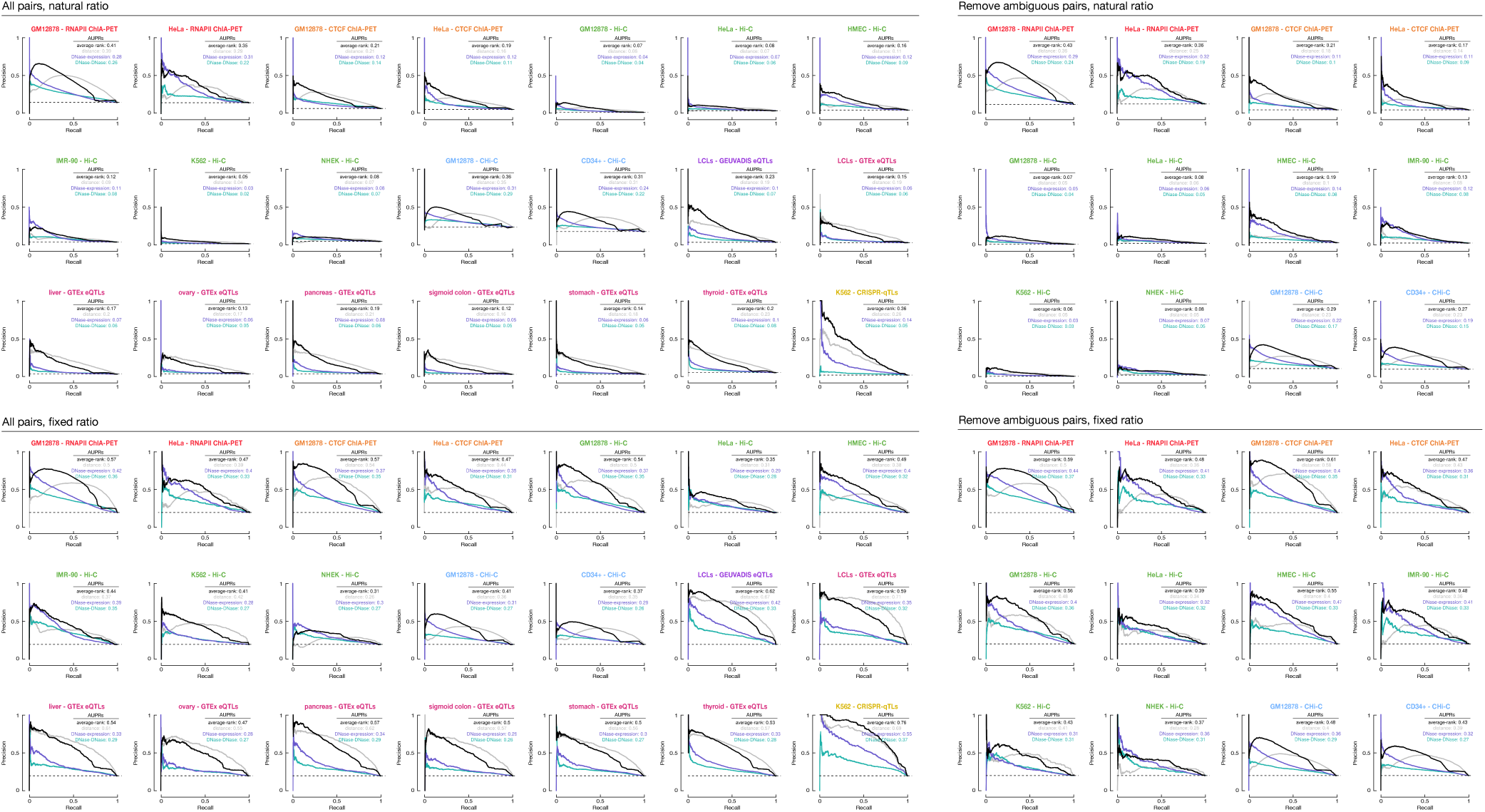
PR curves for unsupervised models. AUPRs for distance (gray), average-rank (black), DNase-DNase correlation (teal), and DNase-expression correlation (purple), across each of the BENGI datasets. Top left group has all pairs with natural ratio. Bottom left group has all paris with fixed ratio. Top right group has ambiguous pairs removed with natural ratio. Bottom right group has ambiguous pairs removed with fixed ratio.

**Supplemental Figure 3.**
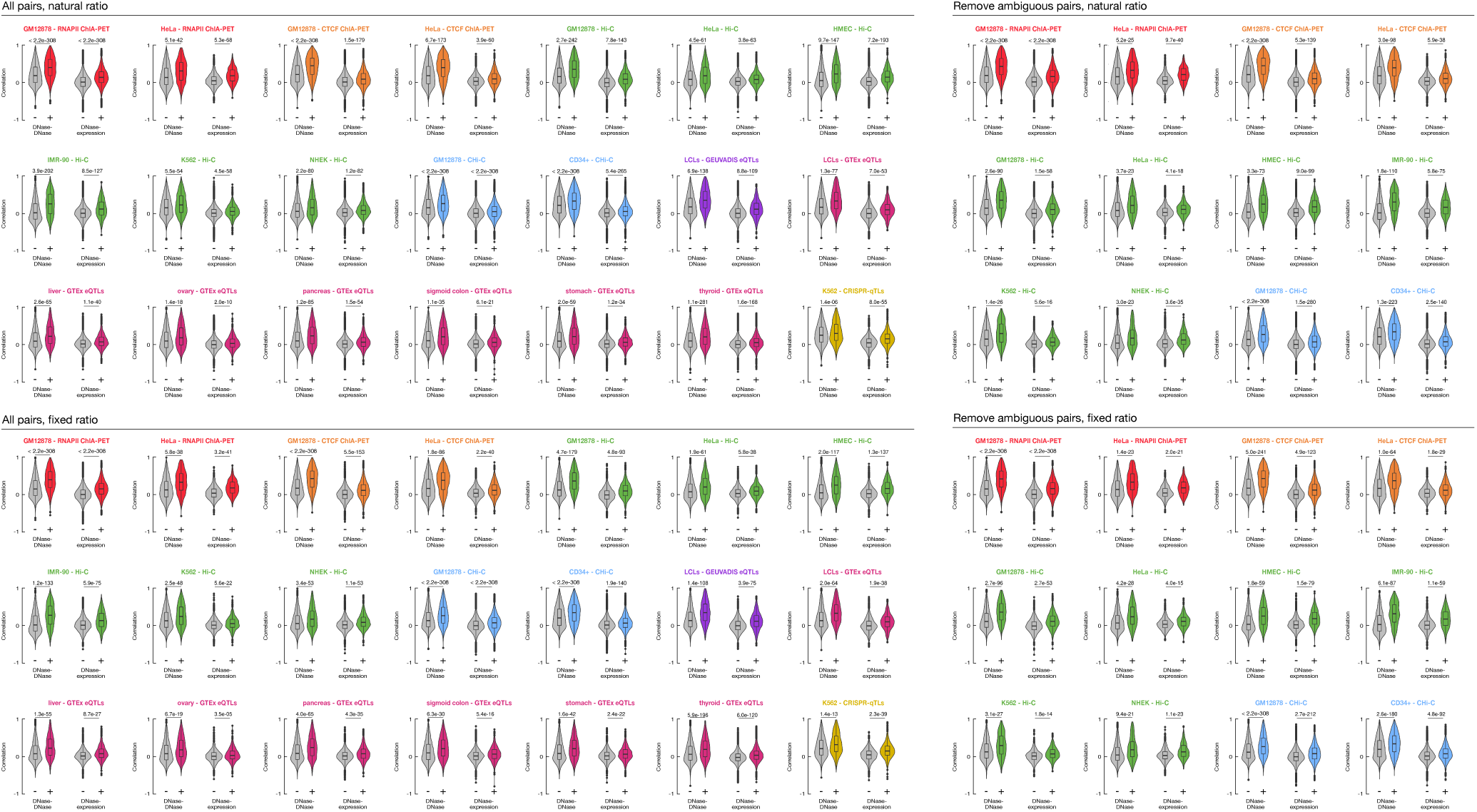
Correlation between BENGI pairs. Violin plots displaying the distributions of Pearson correlation coefficients—computed using the DNase-DNase or DNase-expression method—for positive (right, colored) and negative (left, grey) BENGI pairs. Wilcoxon rank-sum test *p*-values are indicated. Top left group has all pairs with natural ratio. Bottom left group has all paris with fixed ratio. Top right group has ambiguous pairs removed with natural ratio. Bottom right group has ambiguous pairs removed with fixed ratio.

**Supplemental Figure 4.**
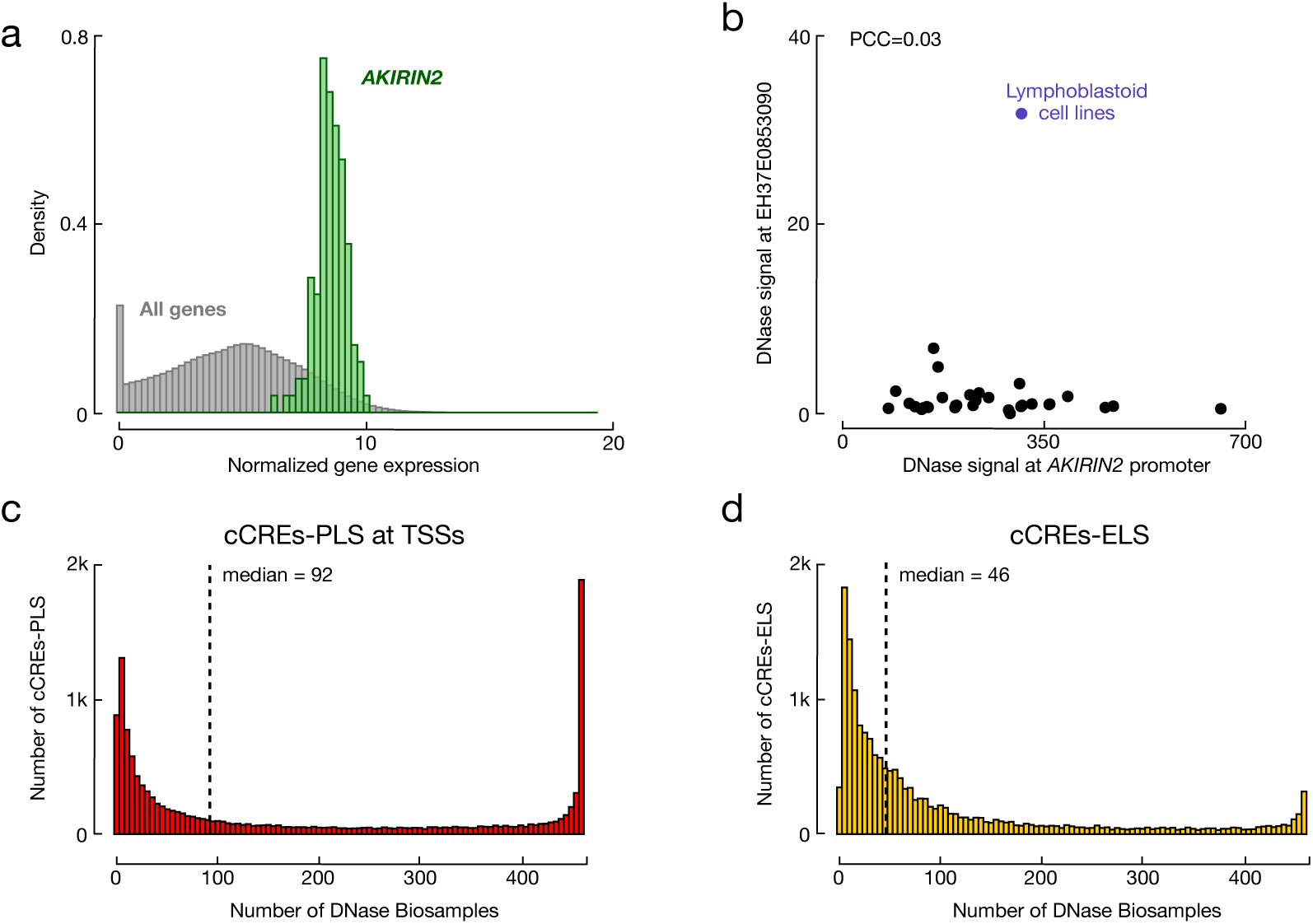
Correlation methods perform poorly due to the ubiquity of promoters. **a**, Normalized gene expression calculated by the DNase-expression method for all genes (black) and *AKIRIN2* (green) across 112 cell types. **b**, DNase signal at EH37E0853090 and *AKIRIN2’s* promoter using the DNase-DNase correlation method. Only the Lymphoblastoid cell line group (purple) has high signal at EH37E0853090. **c-d**, Number of biosamples in the ENCODE phase 2, ENCODE phase 3, and Roadmap projects with high DNase (Z-score > 1.64) for cCREs-TSS with promoter like signatures (cCRE-PLS) and cCREs with enhancer-like signatures (cCREs-ELS) included in BENGI datasets.

**Supplemental Figure 5.**
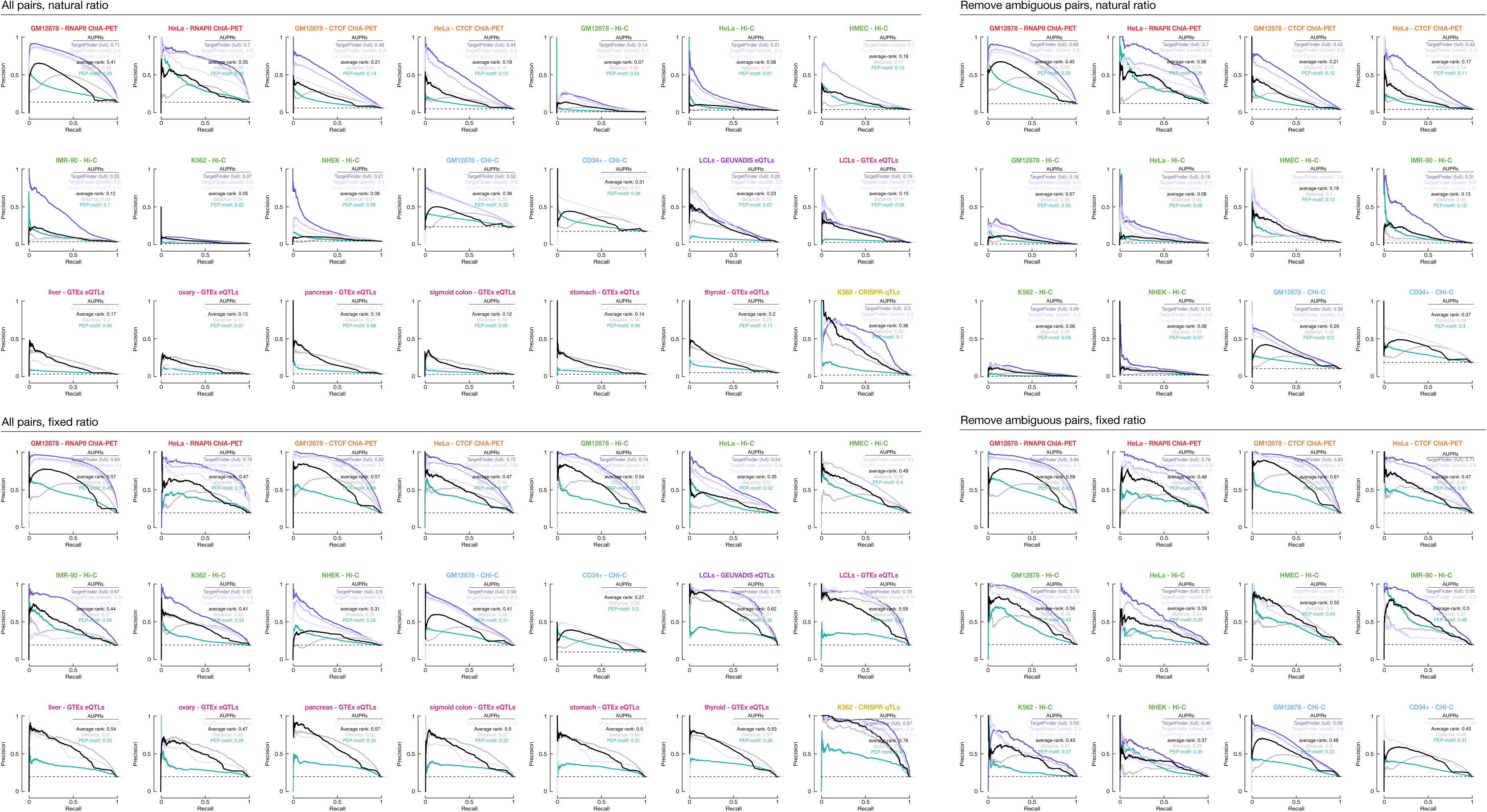
PR curves of supervised methods evaluated on BENGI datasets. AUPRs for distance (gray), average-rank (black), PEP-motif (teal), TargetFinder full-model (dark purple), TargetFinder core4 (medium purple) and TargetFinder core3 (light purple) methods in each of the BENGI datasets. Top left group has all pairs with natural ratio. Bottom left group has all paris with fixed ratio. Top right group has ambiguous pairs removed with natural ratio. Bottom right group has ambiguous pairs removed with fixed ratio.

## TABLES

**Table 1 | Genomic interaction datasets**

**Table 2 | Computation methods for predicting target genes**

## SUPPLEMENTAL TABLES

**Supplemental Table 1: Input data sources**

**Supplemental Table 2: Summary of BENGI datasets**

**Supplemental Table 3: AUPRs for unsupervised methods**

**Supplemental Table 4: AUPR for supervised methods**

**Supplemental Table 5: AUPR for cross-cell type methods**

## METHODS

### Data acquisition

#### ChIA-PET

We downloaded the following ChIA-PET clusters generated by the Ruan lab^31^ from NCBI’s Gene Expression Omnibus (GEO) under the accession GSE72816.

~~~
      GSM1872886_GM12878_CTCF_PET_clusters.txt
      GSM1872887_GM12878_RNAPII_PET_clusters.txt
      GSM1872888_HeLa_CTCF_PET_clusters.txt
      GSM1872889_HeLa_RNAPII_PET_clusters.txt
~~~

We filtered each set of clusters by selecting ChIA-PET links that were supported by at least four reads (column 7 ≥ 4).

#### Hi-C Loops

We downloaded the following Hi-C loops generated by the Aiden lab^28^ from GEO under the accession GSE63525.

~~~
    GSE63525_GM12878_primary+replicate_HiCCUPS_looplist.txt
    GSE63525_HMEC_HiCCUPS_looplist.txt.gz
    GSE63525_HeLa_HiCCUPS_looplist.txt.gz
    GSE63525_IMR90_HiCCUPS_looplist.txt.gz
    GSE63525_K562_HiCCUPS_looplist.txt.gz
    GSE63525_NHEK_HiCCUPS_looplist.txt.gz
~~~

We did not perform any additional filtering on these loops.

#### CHi-C

We downloaded the following CHi-C interactions generated by the Osborne lab^32^ from ArrayExpress under the accession E-MTAB-2323.

~~~
    TS5_GM12878_promoter-other_significant_interactions.txt
    TS5_CD34_promoter-other_significant_interactions.txt
~~~

We filtered each set of interactions selecting CHi-C links by requiring log(observed/expected) greater than ten (column 11 > 10).

#### eQTLs

We downloaded eQTLs from the GEUVADIS project:

~~~
    ftp://ftp.ebi.ac.uk/pub/databases/microarray/data/experiment/GEUV/E-GEUV-1/analysis_results/
    EUR373.gene.cis.FDR5.all.rs137.txt
~~~

We downloaded GTEx eQTLs in `GTEx_Analysis_v7_eQTL.tar.gz` from the gTEX Portal https://gtexportal.org/home/datasets. We used the following files:

~~~
    Cells_EBV-transformed_lymphocytes.v7.signif_variant_gene_pairs.txt
    Colon_Sigmoid.v7.signif_variant_gene_pairs.txt
    Liver.v7.signif_variant_gene_pairs.txt
    Ovary.v7.signif_variant_gene_pairs.txt
    Pancreas.v7.signif_variant_gene_pairs.txt
    Stomach.v7.signif_variant_gene_pairs.txt
    Thyroid.v7.signif_variant_gene_pairs.txt
~~~

#### CRISPR Perturbations

We downloaded crisprQTL data from Gasperini *et al.*^17^ and mapped the reported genes to those annotated in GENCODE V19 and intersected the reported enhancer coordinates with cCREs-ELS in K562. 4,937 of the tested enhancers (85%) overlapped a K562 cCRE-ELS.

### Defining cCREs-ELS

We used cCREs-ELS from V1 of the ENCODE Registry of cCREs available on the ENCODE portal found under the accessions provided in **Supplemental Table 1a**. We selected all cCREs-ELS (RGB color code 255,205,0) that were distal (i.e., greater than 2 kb from an annotated TSS, GENCODE v19).

### Defining cCRE-gene pairs

We created cCRE-gene pairs using the script *Generate-Benchmark.sh*. Which is available on GitHub.

#### 3D chromatin interactions (ChIA-PET, Hi-C, and CHi-C)

Using bedtools intersect (v2.27.1), we intersected the anchors of the filtered links (see above) with cCREs-ELS that were active in the same biosample. We retained all links with an anchor that overlapped at least one cCREs-ELS and with the other anchor within ± 2 kb of a GENCODE V19 TSS. We tagged all links with an anchor within ± 2 kb of the TSSs of multiple genes as ambiguous pairs and created a separate version of each dataset with these links removed.

#### Genetic interactions (eQTLs)

For eQTLs, we retrieved the location of each reported SNP from the eQTL file and intersected these loci with cCRE-ELS that were active in the same tissue type using bedtools intersect. We then paired the cCRE-ELS with the gene linked to the SNP. We only considered SNPs that were directly reported in each of the studies; we did not expand our set using linkage disequilibrium due to the mixed populations surveyed by GTEx.

#### CRISPR/dCas-9 (crisprQTLs)

For crisprQTLs, we intersected the reported positive enhancers with cCREs in K562 using bedtools intersect. We then paired the cCRE-ELS with the gene linked to the reported enhancer.

#### Generating negative pairs

To generate negative pairs, we calculated the 95^th^ percentile of the distances of positive cCRE-gene pairs for each dataset, with distance defined as the linear distance between the cCRE-ELS and the closest TSS of the gene using bedtools closest. For each cCRE-ELS in the positive cCRE-gene pairs that fell within this 95^th^ percentile, we considered all other genes within the 95^th^ percentile distance cutoff as negatives. For datasets with ambiguous links removed (ChIA-PET, Hi-C, and CHi-C), we also excluded genes in these ambiguous pairs as negatives. For the fixed ratio datasets, we also excluded genes that were in the positive pairs for the cCREs-ELS in other BENGI datasets before randomly selecting the negatives. If a cCRE-ELS had fewer than four negative pairs, then it was excluded from this fixed-ratio set.

#### Assigning chromosome CV

For each BENGI dataset, we calculated the number of cCRE-gene pairs on each chromosome and assigned chromCV groups accordingly. The chromosome with the most pairs (often chr1) was assigned its own group. Then, we iteratively took the chromosome with the most and fewest pairs and combined them to create one CV group. In total, the 23 chromosomes (1-22, X) were assigned to 12 CV groups.

### Characterizing BENGI datasets

#### Clustering of dataset overlap

For each pairwise combination of the GM12878/LCL BENGI datasets, we calculated the overlap coefficient of positive cCRE-gene pairs. Then using *hclust*, we performed hierarchical clustering with default parameters.

#### Gene expression

For biosamples with matching RNA-seq data, we downloaded corresponding RNA-seq data from the ENCODE portal (accessions provided in **Supplemental table 1b, Supplemental Figure 1**). For each gene, we calculated the average TPM between the two experimental replicates. To test if there was a significant difference between BENGI datasets with or without ambiguous pairs, we used a Wilcoxon test.

#### ChIP-seq signals

For cCREs-ELS in each positive pair across GM12878 and LCL BENGI datasets, we calculated the average ChIP-seq signal for 140 transcription factors and DNA binding proteins. We downloaded ChIP-seq signal from the ENCODE portal (accession available in **Supplemental Table 2b**) and used UCSC’s *bigWigAverageOverBed* to calculate the average signal across each cCRE. For each BENGI dataset, we then reported the average signal for all cCREs.

### Implementing cCRE-gene prediction methods

#### *Closest-gene* method

We identified the closest TSS to each cCRE-ELS using bedtools closest and GENCODE V19 TSS annotations. We compared two options: using the full set of GENCODE TSSs (with problematic annotations removed) or using only protein-coding GENCODE TSSs. To evaluate performance, we calculated the overall precision and recall for each BENGI dataset (Script: Closest-Gene-Method.sh).

#### *Distance* method

For each cCRE-gene pair, we calculated the linear distance between the cCRE-ELS and the gene’s nearest TSS. To rank these pairs, we took the inverse (1/distance) and calculated the area under the precision-recall curve (AUPR) using a custom Rscript that uses the PROCR library (Script: Run-Distance-Method.sh).

#### *DNase-DNase* correlation method

We used the same DNase-seq datasets as Thurman *et al.* for their DNase-DNase method. We downloaded these legacy datasets that were generated during ENCODE Phase 2 from the UCSC genome browser. For each cCRE-gene pair, we curated a set of cCREs-TSS by determining the closest cCRE for each TSS of the gene. We then calculated the average DNase signal across the nucleotide positions in the cCRE-ELS and cCRE-TSS for each DNase dataset. For similar cell types, as determined in Thurman *et al*., we averaged the DNase signal among these similar cell types in each of the 32 groups to generate 32 values for each cCRE-ELS and cCRE-TSS. We then calculated the Pearson correlation coefficient (PCC) for each cCRE-ELS and cCRE-TSS pair. If a gene had multiple TSSs, we selected the highest PCC of all the cCRE-ELS and cCRE-TSS comparisons. We ranked predictions by their PCC and calculated the AUPR using the PROCR library (Script: Run-Thurman.sh).

#### *DNase-expression* correlation method

To match the legacy data and normalization methods originally used by^20^ we downloaded normalized counts across 112 cell types for the DNase hypersensitive sites or DHSs (*dhs112_v3.bed*) and genes (*exp112.bed*) from http://big.databio.org/papers/RED/supplement/. We intersected each cCRE-ELS with the DHSs curated by^20^. If a cCRE overlapped more than one DHS, we selected the DHS with the highest signal for the cell type in question (i.e., the DHS with the highest signal in GM12878 for GM12878 cCREs-ELS). For each cCRE-gene pair, we then calculated the Pearson correlation coefficient using the 112 normalized values provided in each matrix. cCRE-gene pairs that did not overlap a DHS or did not have a matching gene in the expression matrix were assigned a score of -100. (Script: Run-Sheffield.sh)

#### PEP-motif

We reimplemented PEP-motif to run on our cCRE-gene pairs with chromCV. Like Yang *et al.*, we calculated motif frequency using FIMO^33^ and the HOCOMOCO database (v11 core,^34^). We also added the ± 4 kb padding to each cCRE-ELS as originally described. We concatenated cross-validation predictions and calculated AUPR using PROCR (Script: Run-PEPMotif.sh).

#### TargetFinder

We reimplement TargetFinder to run on our cCRE-gene pairs with chromCV. For features, we used the identical datasets described in Whalen *et al.* for each cell type. We concatenated cross-validation predictions and calculated AUPR using PROCR (Script: Run-TargetFinder-Full.sh).

#### Cross-cell type performance

To test the cross-cell type performance of TargetFinder, we generated core4 and core3 models for each cell type and then evaluated the models on other cell types. To prevent any overfitting, we assigned the chromCV of the test sets to match those of the training sets.

